# Bone Fracture Detection by Electrical Bioimpedance: First Non-Invasive Measurements in Ex-Vivo Mammalian Femur

**DOI:** 10.1101/622936

**Authors:** A. H. Dell’Osa, A. Concu, F. R. Dobarro, J. C. Felice

## Abstract

The fracture of long bones is one of the pathologies of greater demand of systems of medical emergencies, the method used for the diagnosis, the radiology of X-rays, produces damages to the patients and to the hospitals environment. For these reasons, our group is studying the implementation of a new diagnostic technique for the detection of bone fractures by bioimpedance measurements. To simulate a limb, two phantom of bovine femurs (the one with an entire bone and the other with a sawn bone) were constructed and non-invasive Electrical Impedance Spectroscopy measurements were taken on them in order to identify differences in their respective Cole Cole diagrams. Impedance spectroscopy was performed by a frequency sweep between 1 Hz and 65 kHz at a fixed current of 1 mA. The results obtained show wide differences in the Cole Cole diagrams of both phantoms (entire and fractured bone), especially concerning the real component of the, which latter, around the bones section corresponding to that of the lesion in both femurs, was always lower in the fractured femur than the entire one. These first superficial (non-invasive) measurements correspond to the electrical impedance spectroscopy bases and these could -in turn- correspond to what occurs in mammals immediately after the fracture happens, i. e. a dramatic increase in electrical conductivity due to diffusion into the fracture site of more conductive materials such as the blood and the extravascular fluids.

## I. Introduction

Bioimpedance is applied as a medical diagnostic method at the respiratory[1], cardiocirculatory[2], dermatological [3] muscular level[4] and even for the detection of some types of cancer[5].

The suspicion of bone fracture in upper and lower limbs is the main traumatic cause of admission to the public system of medical emergencies in Argentina, the third after parturitions and infections[6]. The diagnostic gold-standart method is X-ray radiology, a technique that is not safe for the patient or for the healthcare environment. The possibility of replacing this classical method with a technique based on bioimpedance measurements on the patient which would generate attention in clinical emergency structures due to less consequences for the patient and the environment, as well as a potential diagnostic method of lower energy consumption compared to a equipment of X-ray emission[7]–[9].

The technique of Electrical Impedance Spectroscopy (EIS) is a typical bioimpedance measurement in which a sinusoidal electrical signal is applied -varying its repetition period or frequency- on a biological system to be studied and thus to know the behavior of the biological system in a defined range of frequencies[10]. The Cole Cole diagram[11] is a classical bioimpedance plot in which the impedance data obtained are presented, corresponding the resistive part with the reactive part of the impedance.

As far as we know, at present there aren’t published papers concerning superficial (non-invasive) measurements of bone tissue with the aim of identify a fracture in these structures both on humans or animals[12]. In these experiments, measurements of EIS at constant current (1mA) were made on two phantoms of *ex-vivo* bones excised from two cows coming from the same farm and having the similar age and weight: the one whole and the other billed. The purpose was that of identifying possible differences in their respective Cole Cole diagrams due to the structural difference between them.

## II. Materials and Methods

### A. Phantom construction

To take non-invasive bioimpedance measurements on both fractured and entire bone, we constructed two cylindrical phantoms with a solidified solution of Agar-Agar at a concentration of 5% dissolved in aqueous solution with a concentration of NaCl to 0.9%, in such a way of furnish a medium ionic conductor[10] and, in turn, a mechanical support of the bone contained inside, as is shown in Figure 1.

**Fig. 1.**
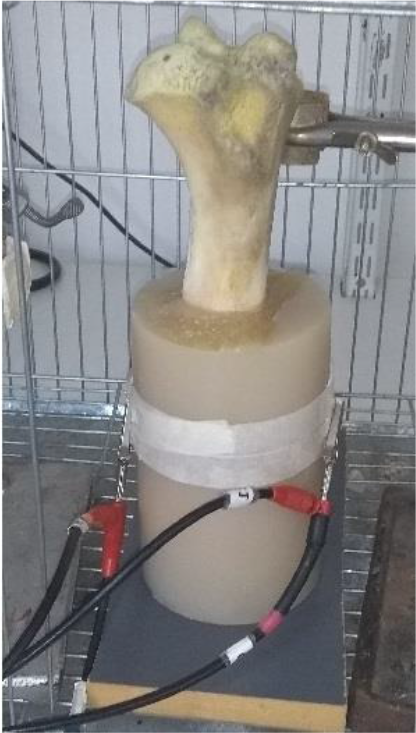
Phantom of entire bone

Figure 2 shows the internal image of both phantoms produced by X-ray radiology.

**Fig. 2.**
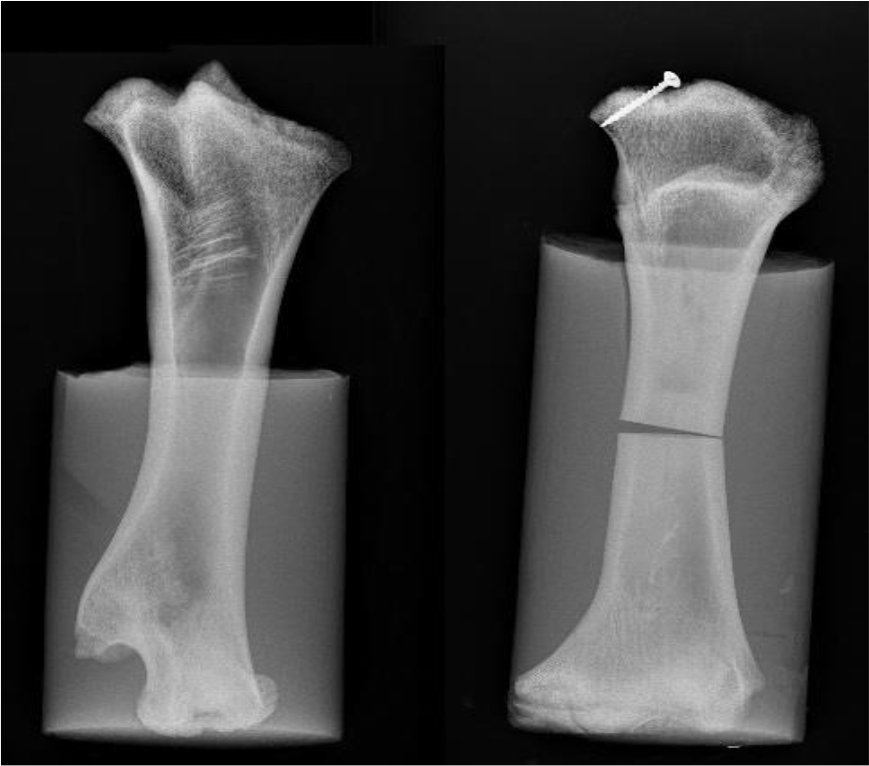
Internal image of both phantoms: on the right radiography is evident the sharp fracture produced by a thin hacksaw. Both femurs show similar height and width.

### B. Measurements

The EIS measurements on the samples were made with a configuration of bipolar electrodes with a Frequency Response Analyzer Solartron 1250/87 (Solartron Metrology Ltd, UK) that injected the 1mA current with a frequency sweep from 1Hz to 65 kHz. The electrodes were disposable electrocardiograph electrodes (3M Ltd, United State) of Ag/AgCl and their diameter was 9 mm. The measurement protocol used was to place the electrodes facing in the axial plane of the phantom, at a certain vertical distance of these respects to the support surface (called height). In each phantom, four EIS measurements were repeated by placing the electrodes couple at a distance from the support surface which was, respectively, of 14, 12, 10 and 8 cm, i.e. an entire space interval that comprised that of the fracture level.

## III. Results

Figure 3 shows the Cole Cole diagram of the measurements, made at the same plane from the base, in both phantoms. From the experimentally assessed data, a circular regression was applied, obtaining characteristic values of the Cole function: R_o_ and R_∞_. Interestingly, experimental data in both phantoms shows a very good compliance with the ideal circumference arch of the corresponding regression. This occurrence reinforces our idea for a good possibility of use the Cole Cole diagram for the bioimpedance studies also in ex vivo bones. Data for the phantom with the entire bone and with the fractured bone for all measurement heights are presented in Table 1.

**Fig. 3.**
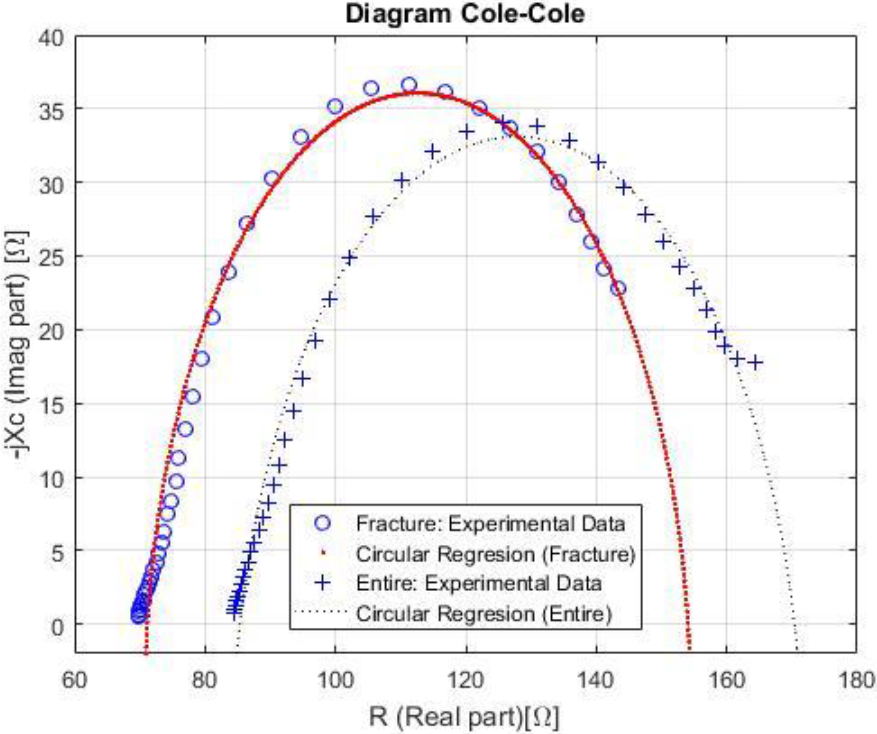
Cole Cole diagram of both phantom with the electrode at 14 cm. The curves drawn with continuous lines represent the circumferential arc calculated by the regression while the points represent the real values.

The Table 1 shows that in the fractured phantom the extreme values of the real part of the impedance were smaller in magnitude than in the whole bone one.

**Table 1.** Values of both R_o_ and R_∞_ for each Cole-Cole diagram

| Height<br>[cm] | Entire bone |  | Fracture bone |  |
| --- | --- | --- | --- | --- |
| | $R_o$ [$\Omega$] | $R_\infty$ [$\Omega$] | $R_o$ [$\Omega$] | $R_\infty$ [$\Omega$] |
| 14 | 170,4 | 85,4 | 154,0 | 71,2 |
| 12 | 162,2 | 79,6 | 146,6 | 68,5 |
| 10 | 172,7 | 82,8 | 162,0 | 75,0 |
| 8 | 181,8 | 95,0 | 167,0 | 72,0 |

## IV. Conclusions

The most important observation in sight, from the results, is obtained with the Cole Cole diagram which is the representative graph of any measurement of EIS on a biological system[10], [13]. Surface measurements - regardless of the height at which the electrodes are located- show a total decrease in the resistive part of the phantom impedance that the bone fracture condition presents with respect to the entire condition. This could be explained because the cross section of the osteotomy in the bone is occupied by a substance with high conductivity (saline solution). Even if in a simplified way, this experimental condition might simulate what happens in a mammalian bone fracture. In fact, when the fracture occurs (no matter what kind of fracture it is), in its proximal tissues there is an increase in blood flow with localized oedema due to the inflammatory reaction and to the process of healing and healing itself bone reconstruction. All of these occurrences imply a diffusion of material with low electrical resistance inside the fracture section, i.e. a mirror condition like that simulated in our fractured bone phantom. Moreover, the relatively large resistance reduction, observed along the longitudinal axis of the sawn bone (about 6 cm), with respect to the entire bone, could be due to the considerable porosity of the trabecular structure of the bone contained in the phantom, a condition, this, that can not be kept very unlike what could happen around an in vivo bone immediately after a fracture.

The limitations present in this work are that measurements were taken on only two phantoms in two different conditions (entire bone and fractured bone), which doesn’t allow having a number of experimental samples to have a determining conclusion. Neither the bones belong to the same animal although both are femur of adult and very similarly structured cow; the comparison between fractured and entire bone would have been ideal to be from the same animal to have an anatomy with greater similarity.

Moreover, due to the needs of mechanically secure together both portion of the fractured bone, we extended the agar gel in that phantom. In a next measurement, both phantoms should be constructed with the bones completely submerged in the agar solution, equaling the volume and the height in both cases, to confirm that, due to the very high conductivity of the agar gel, dispersion of currents (which in the present work we can’t evaluate) prove to be negligible if the Agar gel volume is diverse among phantoms.

At present, there is no bioimpedance or EIS analysis with non-invasive surface electrodes (invasive surgical pins in bone tissue and limb had been used) on living beings to correlate bioimpedance values with bone limbs integrity [12], though attempts of application of Electrical Impedance Tomography (EIT) [14] to generate a diagnostic image has been made, but these had not reached a quality to make a diagnosis. So this work could represent the first approach, from a simplified model such as phantoms built, to take surface measurements; obtaining coherent data with respect to the basic concepts of the EIS (Cole Cole diagram) and compatible with the physiopathological reality of bone fracture (increase in conductivity from the absence of a tissue of greater resistivity as it is the bone that is replaced by blood and extravascular/high conductive fluids) [7].

Reasonably, this work gives the first indications of feasibility to specify the detection of a fracture of a long bone in a limb in human beings through bioimpedance measurements, just strictly after the accident happens. Even though it is a quick analysis of these measurements in which the precise height of the fracture and the geometric measurements of the bones, among other factors to be analyzed, are not contemplated, any way this does not detract this proposed first conclusion from this preliminary analysis.

## Acknowledgment

The authors thank Dr. Martín Zamora, LAMEIN, Argentina, for his collaboration in the development of phantoms, and Dr. Andrea Fois, CEO of the Nomadyca Ltd, Italy, for his contribution in the analysis of the experimental data.

The first author was partially supported by a Grant of the Ministero degli Affari Esteri e della Cooperazione Internazionale (MAECI) of Italy, CONICET of República Argentina, IDEI Universidad Nacional de Tierra del Fuego and Gobierno de la Provincia de Tierra del Fuego. He thanks LAMEIN Argentina, 2C TECHNOLOGIES S.R.L. and Università di Cagliari for the kind hospitality where part of this work has been done.

## References

1. F. Simini, E. Santos, and M. Arregui, “Electrical Impedance Tomography to Detect Trends in Pulmonary Oedema,” in Bioimpedance in Biomedical Applications and Research, 1st ed., F. Simini and P. Bertemes-Filho, Eds. Springer International Publishing, 2017, pp. 45–64.

2. A. Crisafulli, S. Salis, F. Tocco, F. Melis, R. Milia, G. Pittau, M. A. Caria, R. Solinas, L. Meloni, P. Pagliaro, and A. Concu, “Impaired central hemodynamic response and exaggerated vasoconstriction during muscle metaboreflex activation in heart failure patients,” Am. J. Physiol. (Heart Circ. Physiol.), vol. 292, pp. 2988–2996, 2007.

3. R. P. Braun, J. Mangana, S. Goldinger, L. French, R. Dummer, and A. A. Marghoob, “Electrical Impedance Spectroscopy in Skin Cancer Diagnosis,” Dermatol. Clin., vol. 35, no. 4, pp. 489–493, 2017.

4. L. Nescolarde, J. Yanguas, J. Terricabras, H. Lukaski, X. Alomar, J. Rosell-Ferrer, and G. Rodas, “Detection of muscle gap by L-BIA in muscle injuries : clinical prognosis,” Physiol. Meas., vol. 21, no. 38(7), pp. L1–L9, 2017.

5. P. Bertemes-Filho and F. Simini, Bioimpedance in Biomedical Applications and Research. Montevideo, Uruguay: Springer International Publishing, 2018.

6. Dirección de Estadísticas e Información de Salud - Ministerio de Salud de la República Argetina, “Egresos hospitalarios del sector oficial, según edad por grupos de diagnósticos | Deis,” 2014. http://www.deis.msal.gov.ar/index.php/causas-egresos/.

7. Robbins, R. S. Cotran, V. Kumar, and C. T., Patologia estructural y funcional, 6th ed. España: Interamericana, 2000.

8. R. McRae, Tratamiento pr ctico de fracturas, 5th ed. Barcelona, España: Elsevier Health Sciences Spain, 2010.

9. J. G. Webster, E. R. Ritenour, T. Slavik, and N. Kwan-Hoong, Webb’s physics of Medical Imaging, 2nd ed. New York, USA: CRC Press, 2011.

10. M. Valentinuzzi, J. P. Morucci, B. Riguad, and C. J. Felice, “Critical Reviews in Biomedical Engineering,” p. Volume 24, Issues 4-6, 1996.

11. K. S. Cole and R. H. Cole, “Dispersion and absorption in dielectrics I. Alternating current characteristics,” J. Chem. Phys., vol. 9, no. 4, pp. 341–351, 1941.

12. A. H. Dell’Osa, F. Simini, and C. J. Felice, “Bioimpedance and bone fracture detection : A state of the art,” in III Latin-American Conference on Bioimpedance, Manizales-Caldas, Colombia, October 3rd - 5th, 2018.

13. S. Grimnes and Ø. Martinsen, Bioimpedance and bioelectricity basics, 3rd ed. Oslo, Noruega: Academic Publics, 2014.

14. H. C. Jongschaap, R. Wytch, J. M. Hutchison, and V. Kulkarni, “Electrical impedance tomography: a review of current literature.,” Eur. J. Radiol., vol. 18, no. 3, pp. 165–74, 1994.

